# Multisensory brain processing is associated with depressive symptoms in youth with chronic pain

**DOI:** 10.64898/2026.11.14.766198

**Authors:** Laura Martín-Herrero, Maria Suñol, Saül Pascual-Diaz, Tracy V. Ting, Jonathan A. Dudley, Catherine Jackson, Susmita Kashikar-Zuck, Robert C. Coghill, Marina López-Solà

**Affiliations:** Institute of Neurosciences, University of Barcelona, Barcelona, Spain; Department of Medicine, School of Medicine and Health Sciences, University of Barcelona, Barcelona, Spain; IDIBAPS, Institut d’Investigacions Biomèdiques August Pi i Sunyer, Barcelona, Spain; Center for Neuroscience Imaging Research, Institute for Basic Science (IBS), Suwon, South Korea; Department of Pediatrics, University of Cincinnati College of Medicine, Cincinnati, OH, USA; Division of Rheumatology, Cincinnati Children’s Hospital Medical Center, Cincinnati, OH, USA; Division of Behavioral Medicine and Clinical Psychology, Cincinnati Children’s Hospital Medical Center, Cincinnati, OH, USA; Pediatric Pain Research Center, Cincinnati Children’s Hospital Medical Center, Cincinnati, OH, USA; Serra Hunter Programme, Department of Medicine, School of Medicine and Health Sciences, University of Barcelona, Barcelona, Spain

**Keywords:** Adolescence, Chronic Pain, Depressive Symptoms, Functional magnetic resonance (fMRI), Neuroimaging, Sensory Sensitivities

## Abstract

Juvenile fibromyalgia (JFM) is a chronic pain condition marked by widespread musculoskeletal pain and often accompanied by physical fatigue, nonrestorative sleep, headaches, anxiety, and depression. Previous studies have found that adolescents with fibromyalgia report augmented unpleasantness and altered brain responses to non-painful multisensory stimuli. Likewise, research in major depression has shown altered neural processing of sensory stimulation. We examined whether augmented brain responses to non-painful multisensory stimulation is associated with depressive symptoms in adolescent girls with JFM. Forty-six female adolescents with JFM (mean ± SD, 16.43 ± 1.10 years [range 13-18 years]) and 44 healthy participants (mean ± SD, 16.09 ± 1.06 years [range 13-18 years]) completed validated questionnaires assessing daily-life sensory hypersensitivity and depressive symptoms, alongside undergoing a functional MRI multisensory task. JFM patients reported significantly higher sensory unpleasantness than healthy participants, and this hypersensitivity was positively associated with depressive symptoms. Neuroimaging results in chronic pain patients revealed that greater depressive symptomatology was associated with increased activation in the posterior cingulate cortex (PCC) (p_FWE_<0.05), a region implicated in self-referential processing and specifically rumination in adults with chronic pain. JFM patients showed increased PCC-left middle temporal gyrus task-dependent connectivity using psychophysiological interactions analysis (PPI). Furthermore, hyperconnectivity between the PCC and brain areas involved in cognitive-affective regulation, sensory integration, and memory processing was strongly correlated with depressive symptoms in JFM patients. These findings underscore the role of depressive symptoms in sensory processing and integration in JFM patients and highlight the complex interplay between sensory processing and emotional/self-referential brain function in patients.

## 1. Introduction

Chronic pain during adolescence is shaped by psychosocial and environmental factors, including family influences such as parental catastrophizing and sociocultural variables like socioeconomic status and perceived social support, which increase vulnerability to chronic pain and affective symptoms, specifically, depression and anxiety, key predictors of higher pain and functional disability (1–8).

In this context, Juvenile fibromyalgia (JFM) is a paradigmatic example. JFM is a chronic pain syndrome primarily affecting female adolescents characterized by widespread musculoskeletal pain, fatigue, nonrestorative sleep, headaches, anxiety, and depression (9–13). Affective symptoms are particularly significant in JFM, coinciding with maturation of emotion regulation systems and significant social/academic stressors (6,8,14). The interplay between affective dysregulation and chronic pain can create a cycle in which negative mood states intensify pain, undermine coping, and contribute to functional disability (15). In JFM depressive symptoms are associated with greater pain severity and functional impairment, poorer sleep, lower quality of life and predict long-term physical and psychological outcomes (12,14,16).

Neuroimaging studies in adult patients with chronic pain have shown structural and functional alterations in regions involved in emotional and nociceptive processing, including the cingulate cortex, insula, amygdala, and hippocampus (17–22). The posterior cingulate cortex (PCC), a core hub for emotional, cognitive and sensory integration (23), has been associated with both chronic pain and depression (24–26). In chronic pain, altered PCC activity and connectivity have been linked to disrupted default mode network functioning and self-referential pain processing (17,27,28). Similarly, in major depression, PCC dysfunction has been associated with maladaptive self-referential processing and depressive symptomatology (24,26,29).

In addition to alterations in affective and attentional brain networks, sensory processing is disrupted in adults with fibromyalgia (FM) and with depression. Neuroimaging studies showed altered neural responses to sensory stimulation in regions involved in somatosensory integration, salience and affective processing (19,20,30). Similar alterations have been observed in adolescents with JFM and with depression (31–33). These neural findings align with behavioral reports of multisensory hypersensitivity across fibromyalgia and depression across adults and adolescents (19,20,33–36).

Here, we aim to investigate, for the first time, whether depressive symptoms are associated with altered brain responses to non-painful multisensory stimulation in adolescents with JFM. We hypothesized that greater depressive symptoms would relate to alterations in brain function during multisensory processing, particularly in regions involved in sensory integration, affective processing, and self-referential functioning, such as the PCC and sensory integration networks. We also examined healthy adolescent girls to assess group differences in task-evoked brain connectivity within regions associated with greater depressive symptomatology.

## 2. Methods

### 2.1 Participants

Ninety female adolescents participated in the study, including 46 JFM patients (mean ± SD, 16.43 ± 1.10 years [range 13-18 years]) and 44 healthy participants (mean ± SD, 16.09 ± 1.06 years [range 13-18 years]) (**Table 1**). Participants with JFM were recruited from the parent clinical trial, the Fibromyalgia Integrative Training for Teens (FIT-teens) study (37). This dataset has been previously used to examine other research questions (32,33); however, the present study investigates a novel question and outcome not addressed in prior reports. To ensure transparency and minimize concerns regarding type I error inflation across studies using this dataset, the study design, sample size calculations, and analysis plan were preregistered on the Open Science Framework (OSF) website (Registration DOI: https://doi.org/10.17605/OSF.IO/4ET5F) (38). Ethical approval was obtained from the Institutional Review Board of Cincinnati Children’s Hospital Medical Center (2017–7771), and informed assent and parental consent were obtained. See **Supplementary Methods 1** for further details.

**Table 1.** Demographic and clinical characteristics of the study sample.

|  | Healthy Participants (n=44) | JFM Patients (n= 46) | Statistics |  |
|---|---|---|---|---|
| DEMOGRAPHIC VARIABLES | Mean ± SD | Mean ± SD | T / X2 | p-value |
| Age (years) | 16.09 ±1.06 | 16.43 ±1.10 | 1.51 | 0.13 |
| Race (C / NC) | 2.06 | 2.1 | 0.54 | 0.58 |
| Yearly Household Income (1–7) | 5.18 ±1.90 | 4.86 ±1.96 | -0.76 | 0.44 |
| Education Level of the Primary Caregiver (1–5) | 4.41 ±0.86 | 3.95 ±0.89 | -0.38 | 0.39 |
| Education Level of the Secondary Caregiver (1–5) | 3.80 ±0.84 | 3.80 ±1.03 | 0 | 1 |
| **MEDICATION USE** | **no. (%)** | **no. (%)** | **T / X2** | **p-value** |
| Pain-related drugs † | - | 20 (43.47) | - | - |
| Psychiatric drugs‡ | - | 19 (41.30) | - | - |
| Gastrointestinal drugs | - | 5 (10.86) | - | - |
| Antihistamines | - | 2 (4.34) | - | - |
| Melatonin | - | 6 (13.04) | - | - |
| Vitamins/minerals | - | 3 (6.52) | - | - |
| Birth control | 1 (2.27) | 3 (6.52) | - | - |
| Steroids | - | 1 (2.17) | - | - |
| Diuretics | 1 (2.27) | - | - | - |
| Statins | 1 (2.27) | - | - | - |
| **CORE JFM SYMPTOMS** | Mean ± SD | Mean ± SD | T | p-value |
| Functional Disability Inventory | 0.61 ± 1.29 | 23.95 ± 9.05 | 17.30 | <0.0001 |
| Widespread Pain Index | 0.28 ± 1.08 | 11.75± 3.50 | 20.62 | <0.0001 |
| Fibromyalgia Symptom Severity | 1.12 ± 1.40 | 8.93 ± 2.15 | 19.07 | <0.0001 |
| **CHILD DEPRESSION INVENTORY 2** | Mean ± SD | Mean ± SD | T | p-value |
| CDI-2 Total Score | 5.44 ± 5.58 | 19.55 ±10.06 | 7.80 | <0.0001 |
| ◊ CDI-2 Negative Mood | 1.35 ± 1.82 | 8.11 ± 3.75 | 10.50 | <0.0001 |
| ◊ CDI-2 Negative Self Esteem | 0.84 ± 1.15 | 2.95 ± 2.63 | 4.75 | <0.0001 |
| ◊ CDI-2 Ineffectiveness | 2.43 ± 2.46 | 6.46 ± 3.60 | 5.94 | <0.0001 |
| ◊ CDI-2 Interpersonal Problems | 0.46 ± 0.82 | 2.02 ± 1.51 | 5.85 | <0.0001 |
| **SENSORY SENSITIVITY VARIABLES** | Mean ± SD | Mean ± SD | T | p-value |
| AASP Visual | 10.82 ± 3.44 | 16.23 ± 5.34 | 5.50 | <0.0001 |
| AASP Touch | 12.38 ± 3.90 | 21.00± 6.86 | 7.06 | <0.0001 |
| AASP Auditory | 8.51 ± 2.97 | 15.18 ± 5.42 | 6.99 | <0.0001 |

Note: **Yearly household income** is shown using a scale of 1–7, where 1 = <\$24,999; 2 = \$25,000 to \$49,999; 3 = \$50,000to \$74,999; 4 = \$75,000 to \$99,999; 5 = \$100,000 to \$124,999; and 6 = \$125,000 to \$149,999; 7 > \$150,000. **Primary caregiver** refers to the person who is primarily responsible for meeting the child’s daily needs. **Secondary caregiver** refers to the person who also plays a significant role in the child's care and well-being but may not have the primary responsibility. **Caregiver education level** is shown using a scale of 1–5, where 1 = less than high school; 2 = high school/GED;3 = partial college or trade school; 4 = college graduate; 5 = postgraduate degree. AASP, Adult Adolescent Sensory Profile; C, Caucasian; JFM, juvenile fibromyalgia; NC, Non-Caucasian; SD, standard deviation.
<sup>†</sup>Pain-related drugs included antiepileptic drugs, nonsteroidal anti-inflammatory drugs, muscle relaxants, acetaminophen, or acetylsalicylic acid.
<sup>‡</sup>Psychiatric drugs included antidepressants, anxiolytics, and attention deficit hyperactivity disorder drugs.

### 2.2 Clinical measures

Developmentally appropriate and validated self-report measures were used to assess core disease symptoms and functional disability. Participants completed the Pain and Symptom Assessment Tool (PSAT) (36,39), a standardized tool for the classification of fibromyalgia in pediatric patients. PSAT includes the Fibromyalgia Symptom Severity scale (FMSS), measuring FM symptoms experienced in the past week, and the Widespread Pain Index (WPI), which assesses the number of body areas where they had pain during the last three months (13,39). Participants self-reported functional disability (i.e., physical difficulties in daily activities) using the Functional Disability Inventory (FDI) (40).

### 2.3 Affective measures: Child Depression Inventory 2

Developmentally appropriate and validated self-report measures were used to assess depressive symptoms. Participants completed the Child Depression inventory 2 (CDI-2) a 28-item instrument that evaluates depressive symptoms in children and adolescents (41,42). The CDI-2 provides a total score, derived from the sum of all items, as well as four subscale scores assessing distinct symptom dimensions: Negative Mood (e.g., sadness, crying, worry, indecisiveness), Interpersonal Problems (e.g., social difficulties, avoidance, isolation), Negative Self-Esteem (e.g., low self-worth, feeling unloved), and Ineffectiveness (e.g., poor self-evaluation of ability and academic performance).

### 2.4 Daily sensory measures

To assess hypersensitivity to daily non-painful sensory stimuli we used the Adolescent Adult Sensory Profile (AASP) (43). We focused on two subscales of interest: sensory sensitivity and sensory avoidance across 3 modalities of interest (visual, tactile and auditory processing). As is standard, we derived a total score for each modality computed as the addition of the two subscale scores. These subscales have been used in previous adult and adolescent FM research, and effectively capture subjective hypersensitivity to sensory stimuli (20,32,33,44).

### 2.5 Multisensory task description

We used a block design fMRI paradigm involving 4 trials of alternating 30 second periods of rest (no stimulation) and 30 second periods of activation (concurrent visual, auditory, and tactile motor stimulation) (19,20). Activation periods consisted of the simultaneous presentation of visual stimulation (at a temporal frequency of 3 Hz [equivalent to 6 color reversals per second] with a full-field flashing checkerboard composed of a grid of black and white alternating squares [80 ± 10 lux]) and auditory stimulation (a series of 15 tones of frequencies in the range of 233.1–1,318.5 Hz, presented at a temporal frequency of 3 Hz, with an intensity of 75 ± 5 dB), coupled with a finger-opposition task during which the subject was instructed to touch the tip of the right thumb with the other fingers (from index finger to little finger). After each period of multisensory stimulation and a short interstimulus interval ranging from 2-4 seconds, we instructed participant to rate the unpleasantness of the trial (10-second interval), using a computerized visual analogue scale ranging from “not at all unpleasant” (0) to “the most unpleasant you can imagine’’ (100). At the end of the task, participants used visual analogue scales to rate the intensity, unpleasantness, and spread of bodily pain experienced throughout the task, as well as the extent to which bodily pain distracted them.

### 2.6 MRI acquisition parameters and Imaging processing

MRI data acquisition parameters and image preprocessing are detailed in the **Supplementary Methods 2**.

### 2.7 Psychophysiological interactions

Functional connectivity was conducted using psychophysiological interaction (PPI) analysis, implemented in the Statistical Parametric Mapping (SPM) software (SPM12; The Wellcome Department of Imaging Neuroscience). PPI is a statistical approach used to explore task-related changes in connectivity by identifying brain regions whose activity covaries with a seed region during a specific contrast of interest.

#### 2.7.1 Seed region definition

We selected the posterior cingulate cortex (PCC) as the seed region based on a priori hypotheses and prior evidence implicating its role in affective components of pain, including pain-related fear, catastrophizing, avoidance, and rumination. Seed time series were extracted for each participant from the entire PCC mask and from ventral (area 23) and dorsal (area 31) subregions, defined according to the Brainnetome Atlas (Cha et al., 2017; Vogt et al., 2006). Time series were adjusted for the psychological regressor (task condition), the physiological regressor (PCC activity), and their interaction (PPI), using spm_peb_ppi in SPM12.

Supplementary analyses report (i) overlap between PPI seed regions and brain responses associated with depressive symptoms during multisensory stimulation, and (ii) results using a finer PCC parcellation into four subregions (46,47) (**Supplementary Figure 1-2 and Supplementary Table 1**).

### 2.8 Statistical analysis

#### 2.8.1 Behavioral

Preliminary analyses were conducted to inform covariate selection for subsequent models. Two-sample *t*-tests were computed in R software (R Core Team, 2021; R-project.org) to assess between-group differences in core disease features, CDI-2 and daily sensory hypersensitivities AASP, with the aim of informing subsequent covariate selection. We used bivariate Pearson correlations to assess the associations between CDI-2 and daily sensory hypersensitivities AASP. To account for multiple comparisons, Bonferroni correction was applied.

#### 2.8.2 fMRI analyses

In first-level analyses, generalized linear models were implemented in SPM12 to identify task-related activation during multisensory stimulation relative to baseline. Contrast images for the task regressor were entered into second-level analyses to assess the association between brain activation and depressive symptoms (CDI-2). Statistical significance was set at family-wise error (FWE) p_FWE-_<0.05 at the cluster level and p<0.001 at the voxel level. Post-hoc analyses examined associations between brain activation and CDI-2 subscales (Negative Mood, Interpersonal Problems, Negative Self-Esteem, and Ineffectiveness). Further details are provided in the **Supplementary Methods 3**.

#### 2.8.3 PPI analyses

First-level PPI models were computed for each participant using SPM12, including the physiological regressor (seed time series), the psychological regressor (task contrast), and their interaction term (PPI regressor). Nuisance regressors, including head motion parameters and additional confound regressors derived from preprocessing (e.g., FD and spike regressors), were included to control for confounding effects. The PPI individual contrast images were then entered into second-level random-effects analyses. Between-group differences in task-related connectivity (JFM vs. healthy participants) were assessed using two-sample t-test. To investigate the relationship between functional connectivity and depressive symptoms, a second-level GLM was conducted with CDI-2 total score as a covariate. Significance was determined using a voxel-wise threshold of p<0.001 (uncorrected), with cluster-level FWE correction at p<0.05.

In addition, post hoc anatomical labeling of significant PPI clusters was performed using the Brainnetome Atlas (45). We computed the overlap between significant clusters and the atlas to identify their regional composition and provide a fine-grained parcellation of connectivity findings.

Functional decoding was performed using the Neurosynth database to identify cognitive terms most associated with the unthresholded *t*-map of the contrast to preserve the full spatial pattern of brain–behavior associations. A word cloud representing the 15 most strongly correlated functional terms was generated using Python. The size of each term in the word cloud reflects the correlation coefficient between that term and the whole-brain connectivity *t*-map.

## 3. Results

### 3.1 JFM patients with elevated depressive symptoms show higher levels of sensory hypersensitivity in daily life

JFM patients reported significantly higher core JFM symptoms: WPI (t=20.62, P<0.001), FMSS (t=19.07, P<0.001) and FDI (t=17.30, P<0.001) and significantly higher depressive symptoms: CDI-2 total score (t=7.80, P<0.0001), CDI-2 negative mood (t=10.50, P<0.0001), CDI-2 negative self-esteem (t=4.75, P<0.0001), CDI-2 Ineffectiveness (t=5.94, P<0.0001) and CDI-2 Interpersonal Problems (t=5.85, P<0.0001) compared to healthy participants (**Figure 1 and Table 1**). JFM patients also showed significantly higher daily sensory hypersensitivities as reported in the AASP for visual (t=5.50, P<0.0001), tactile (t=7.06, P<0.0001) and auditory (t=6.99, P<0.0001) sensory modalities (**Figure 1 and Table 1**). **Figure 1 and Table 2** show strong positive correlations between patients’ daily life sensory hypersensitivities and depressive symptoms, indicating that patients reporting greater depressive symptoms show higher hypersensitivity (hyper-unpleasantness and behavioral avoidance) to non-nociceptive sensory stimuli of different modalities.

**Figure 1.**
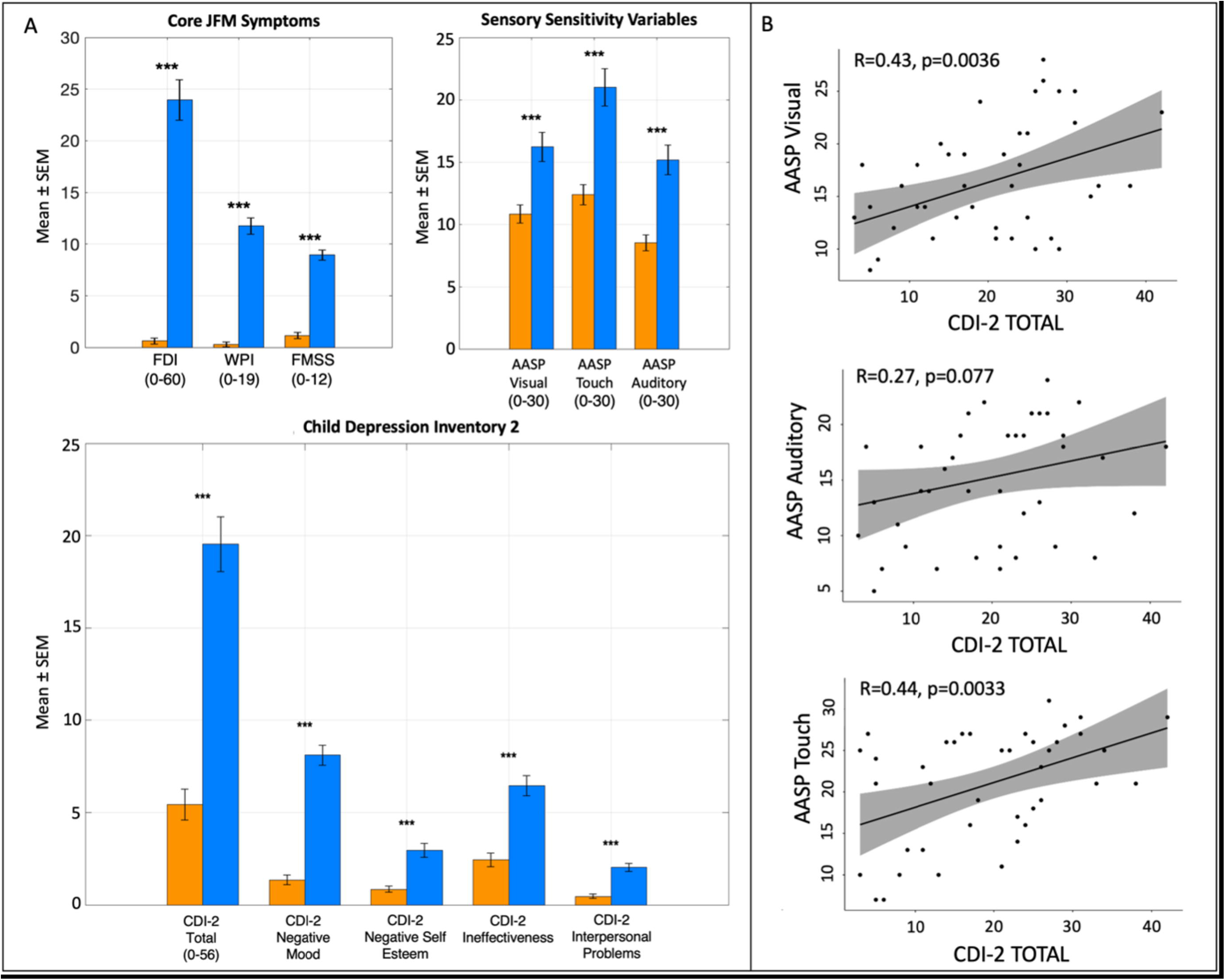
Core symptoms, sensory sensitivity variables, and depressive symptoms in juvenile fibromyalgia (JFM) patients. A) JFM patients showed higher pain-related symptoms (WPI, FMSS, FDI), sensory sensitivity variables (AASP visual, touch and auditory) and depressive symptoms (CDI-2 total and subscales) compared with healthy participants. B) Positive correlations between sensory sensitivity variables (AASP) and Child Depression Inventory 2 Total, indicating that greater depressive symptoms were associated with increased sensory hypersensitivity and avoidance responses to visual, tactile, and auditory stimuli. *P < 0.05, **P < 0.01, ***P < 0.001.

**Table 2.**
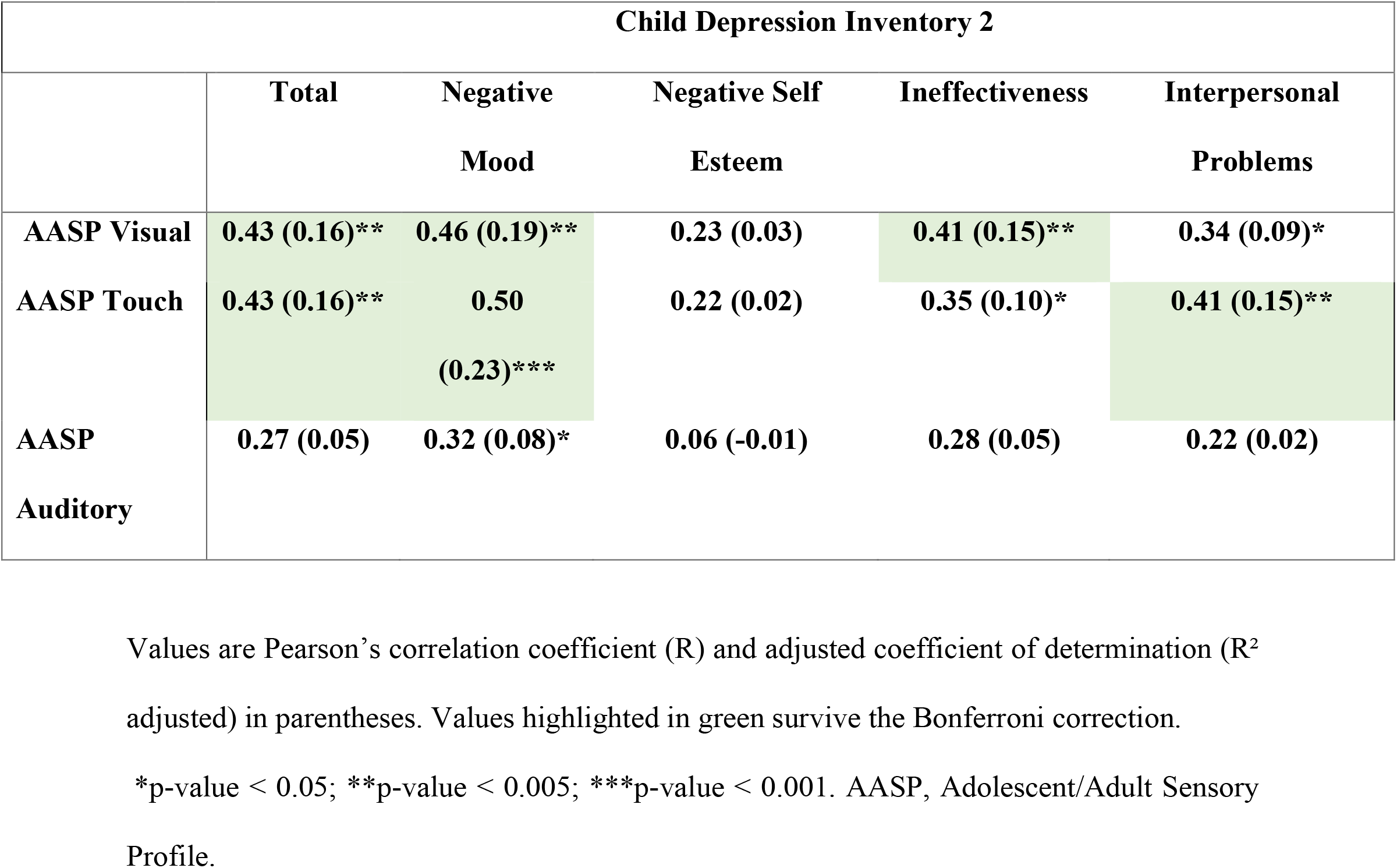
Bivariate correlations between Child Depression Inventory 2 and sensory sensitivities in Juvenile Fibromyalgia patients.

|  | **Child Depression Inventory 2** |  |  |  |  |
|---|---|---|---|---|---|
|  | **Total** | **Negative Mood** | **Negative Self Esteem** | **Ineffectiveness** | **Interpersonal Problems** |
| **AASP Visual** | **0.43 (0.16)\*\*** | **0.46 (0.19)\*\*** | **0.23 (0.03)** | **0.41 (0.15)\*\*** | **0.34 (0.09)\*** |
| **AASP Touch** | **0.43 (0.16)\*\*** | **0.50 (0.23)\*\*\*** | **0.22 (0.02)** | **0.35 (0.10)\*** | **0.41 (0.15)\*\*** |
| **AASP Auditory** | **0.27 (0.05)** | **0.32 (0.08)\*** | **0.06 (-0.01)** | **0.28 (0.05)** | **0.22 (0.02)** |

Values are Pearson’s correlation coefficient (R) and adjusted coefficient of determination (R<sup>2</sup> adjusted) in parentheses. Values highlighted in green survive the Bonferroni correction.
\*p-value < 0.05; \*\*p-value < 0.005; \*\*\*p-value < 0.001. AASP, Adolescent/Adult Sensory Profile.

### 3.2 Depressive symptoms are associated with greater brain responses to multisensory stimulation in JFM patients

In JFM patients, we found whole brain-corrected positive associations between brain activation during the multisensory task and CDI-2 scores, with large effect sizes (Cohen’s d = 1.37–1.52). Specifically, higher total CDI-2 scores, as well as higher scores on the subscales measuring ineffectiveness and negative mood, positively correlated with activation in the right dorsal PCC during multisensory stimulation (FWE cluster-corrected p < 0.05, voxel-level p < 0.001). Additionally, at an exploratory threshold (uncorrected p < 0.005, cluster extent > 20 voxels), scores on the CDI-2 subscale measuring interpersonal problems positively correlated with activation of the right ventral PCC during the task (**Figure 2, Supplementary Table 2**).

**Figure 2.**
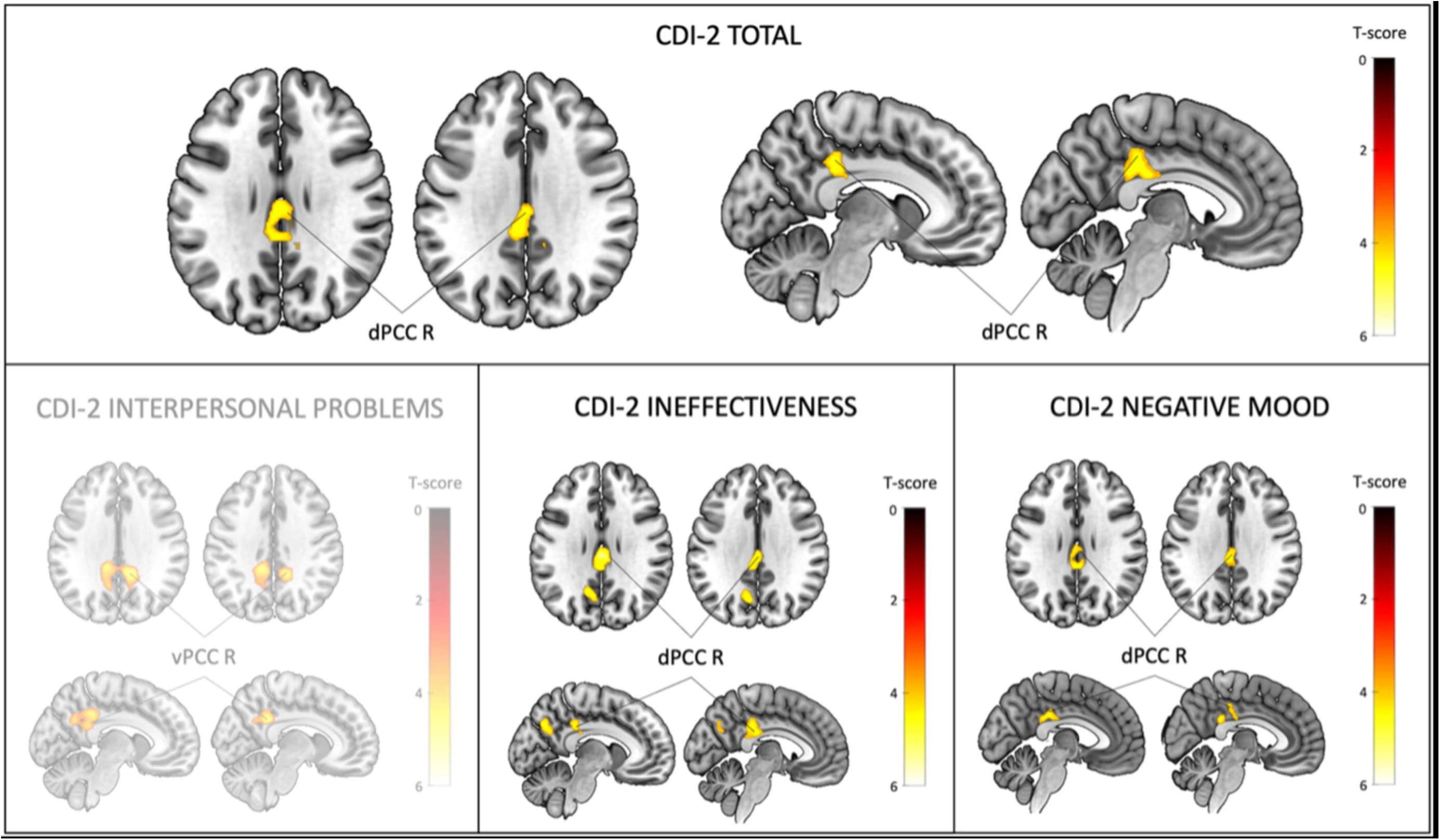
Correlations between task-evoked brain activation and depressive symptoms (CDI-2) in JFM patients. Brain maps show significant correlations between task-evoked brain activation and depressive symptoms. For the total CDI-2 scores and the subscales measuring ineffectiveness and negative mood, statistical significance in second-level analyses was defined as FWE cluster-corrected p < 0.05, with a voxel-level threshold of p < 0.001. For the subscale measuring interpersonal problems, results are presented at an exploratory threshold: voxel-wise p < 0.005 (uncorrected), with a minimum cluster extent of 20 voxels and a cluster-level FWE correction at p < 0.05. CDI-2, Child Depression Inventory 2; d, dorsal; R, right; v, ventral.

At the behavioral level, healthy participants showed a non-significant negative correlation between CDI-2 total scores and right PCC activation (r = –0.10, p = 0.53). The direct comparison between the two correlation coefficients (JFM patients vs. healthy participants) was statistically significant, indicating that the association observed in patients is not present in healthy participants and suggests clinical specificity in the observed effect.

To rule out potential confounding effects of head motion, we first compared mean framewise displacement (FD) between groups and found no significant differences (t = 1.33, p = 0.18). In JFM patients, mean FD was not associated with task-evoked activation (r = –0.09, p = 0.55). Importantly, including FD as a covariate in the second-level analysis did not alter the main results, indicating that head motion did not drive the observed associations. Consistently, no significant effects were observed in healthy controls at either corrected or exploratory thresholds.

### 3.3 Hyperconnectivity of the PCC during multisensory processing in JFM patients is associated with depressive symptoms

Compared to healthy participants, adolescents with JFM exhibited increased task-related PPI connectivity between the PCC region and the left middle temporal gyrus. Notably, this pattern of hyperconnectivity was also observed when using the ventral PCC subregion (area 23) as the seed (voxel-wise p < 0.001, cluster-level FWE-corrected p < 0.05) (**Figure 3, Supplementary Table 3)**.

**Figure 3.**
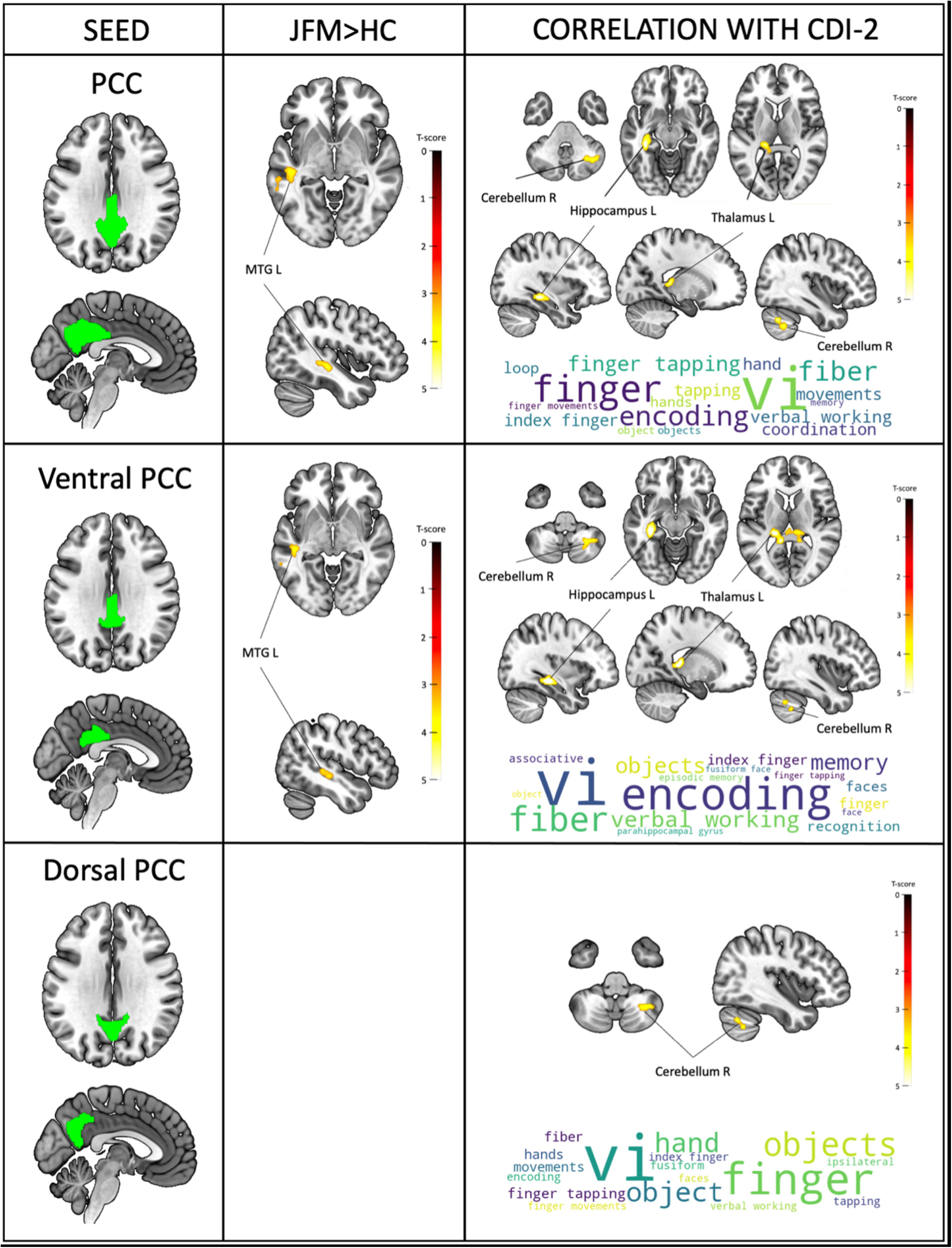
Brain maps showing PPI connectivity differences for each seed of interest (entire PPC region, ventral PCC subregion (Brainnetome area 23) and dorsal PCC subregion (Brainnetome area 31)) between JFM patients and healthy controls and correlations between PPI-based connectivity maps for each seed and depressive symptoms in JFM patients. The bottom right of the figures shows the 15 functional terms most associated with the unthresholded brain t-map of results for each analysis based on the meta-analytic database of Neurosynth (68). Wordcloud was computed with Python. The size of the word is based on the correlation coefficient between each specific term and the connectivity t-map. The color reflects aesthetic purposes. CDI-2, Child Depression Inventory 2. MTG, middle temporal gyrus; L,left; R, right.

In addition, we found positive associations between PCC PPI connectivity and depressive symptoms, specifically in adolescents with JFM. Depressive symptoms, measured using the CDI-2 total score, were positively associated with increased PPI connectivity between the PCC and the left caudal hippocampus, left thalamus, and right cerebellum. Additionally, when using the ventral PCC subregion (Brainnetome area 23) as the seed of interest, we observed higher connectivity with the left caudal hippocampus, left caudal temporal thalamus, and right cerebellum. Finally, using the dorsal PCC subregion (Brainnetome area 31) as the seed of interest, we found increased connectivity only with the right cerebellum (**Figure 3, Supplementary Table 3)**.

To further illustrate the direction of the relationship observed in the voxel-wise PPI regression analyses, we extracted mean connectivity estimates (beta values) from significant clusters identified in the voxel-wise regression analyses and conducted correlations with CDI-2 scores. In JFM patients, mean connectivity estimates extracted from the significant cluster showed a positive but non-significant association with CDI-2 scores (r = 0.21, p = 0.17), whereas healthy participants showed a non-significant negative association (r = –0.16, p = 0.35). The direct comparison between correlation coefficients did not reach statistical significance (p = 0.11). These results are reported descriptively to complement the voxel-wise findings.

To assess the specificity of this effect, the same second-level PPI regression analysis was repeated in healthy controls using CDI-2 scores as a covariate of interest. No significant associations were detected at the same voxel-wise and cluster-level thresholds, suggesting that the connectivity–symptom relationship observed in patients was not present in healthy participants.

To ensure that head motion did not confound the observed connectivity effects, we extracted mean FD for each participant and tested its association with mean beta estimates of task-dependent PPI connectivity extracted from significant clusters in JFM patients. No significant association was observed (r = 0.07, p = 0.62), suggesting that individual differences in head motion were not significantly associated with the observed variability in connectivity measures. In a further sensitivity analysis, mean FD was included as a covariate of no interest in the second-level regression model assessing the association between CDI-2 symptoms and PPI connectivity. The results remained unchanged after controlling for motion, indicating that the observed connectivity–symptom relationships were not driven by head motion.

## 4. Discussion

To our knowledge, this is the first study assessing the relationship between depressive symptoms and hypersensitivity to non-painful sensory stimuli in youth with chronic primary widespread pain using brain imaging. JFM patients reported greater sensory hypersensitivity in daily life and higher depressive symptoms than healthy participants. Furthermore, within the JFM group, higher depressive symptoms were positively associated with greater sensory hypersensitivity. At the brain level, depressive symptoms were associated with increased PCC engagement during multisensory processing, as indicated by significant voxel-wise PPI effects. In contrast, post hoc analyses based on mean connectivity estimates extracted from these clusters showed a non-significant association (r = 0.21), indicating that the voxel-wise effects are not captured by summary-level correlations. These findings suggest that PCC involvement is modulated by individual differences in depression rather than reflecting an amplified response to multisensory stimulation in JFM (32,33). Additionally, compared to healthy participants, JFM patients exhibited PPI hyperconnectivity between the PCC and the left middle temporal gyrus, a region implicated in semantic memory, conceptual processing, and multimodal integration, including auditory and visual information (48–51). Also, during multisensory stimulation, the PCC becomes more tightly functionally coupled with hippocampal, thalamic, and cerebellar structures in individuals with higher depressive symptoms. These findings highlight the central role of the PCC in integrating sensory, cognitive, and affective information, supporting processes related to embodiment, self-reference, memory, and emotion, and providing a neurobiological link between depressive symptoms and altered sensory processing in JFM.

Adolescent females with JFM reported more depressive symptoms than healthy participants. These findings align with previous clinical reports highlighting the high prevalence (22%) of depression among JFM patients (52). Crucially, daily sensory hypersensitivities positively correlated with depressive symptoms in the JFM group, indicating that heightened sensory sensitivity may be linked to greater emotional distress. These findings underscore the importance of investigating the interaction between depressive symptoms and sensory processing in future research to inform clinical assessments and individualized interventions.

As predicted, in JFM patients, depressive symptoms were associated with increased activation in the PCC, during non-painful multisensory stimulation. Importantly, this engagement was not observed as a general task effect, but emerged specifically when individual differences in depressive symptom severity were taken into account, consistent with our previous multisensory neuroimaging work in youth with JFM, in which disease severity is most strongly associated with brain function abnormalities (32,33). Previous studies have implicated the posterior cingulate cortex (PCC) as a key hub within large-scale brain networks involved in pain processing and self-referential cognition, supporting the interaction between sensory-discriminative and affective–cognitive dimensions of pain, as well as its modulation in chronic pain conditions (53,54). Building on this evidence, we emphasize the relevance of PCC activity in evaluating multisensory inputs and integrating them with emotional, self-relevant context to shape cognitive-affective responses to such input. This integration is especially relevant in chronic pain and depression, two conditions that involve altered perception and affective appraisal of sensory stimuli beyond nociceptive signals (15). Consistent with this view, alterations in PCC connectivity have been linked to depression and a higher risk of suffering depression in healthy population, specifically in young adults (26).

At a more fine-grained level, we observed strong associations (Cohen’s d > 1) between increased brain responses to multisensory stimulation in the right dorsal PCC (dPCC) and higher total depressive symptoms (i.e., CDI-2 total scores), as well as the CDI-2 ineffectiveness and negative mood subscales. In contrast, increased activation in the right ventral PCC (vPCC) was strongly associated with higher scores on the CDI-2 interpersonal problems subscale. The dPCC is functionally positioned between the posterior DMN and networks supporting cognitive and emotional control, facilitating attentional allocation and dynamic switching between internal and external focus environments (23,55). Heightened dPCC engagement may therefore reflect increased internal monitoring and dysregulated cognitive–affective regulation, potentially associated with augmented negative self-referential thought and mood disturbances, as previously observed in depressive disorders (23,56–58). The vPCC is more specifically involved in internally directed cognition, including autobiographical memory and social– emotional processing (23,56–58), suggesting that its over-engagement in patients with higher depression symptoms may be associated with ruminative reflection on negative interpersonal experiences and social withdrawal. Together, these findings suggest an important role of the PCC in mediating the complex interaction between sensory processing, embodiment, memory, self-reference, and emotion in JFM. Engagement of the PCC during non-painful multisensory stimulation may reflect altered cognitive–affective appraisal of sensory input, providing insight into shared neurobiological mechanisms underlying chronic pain and depressive symptoms.

Functional connectivity of the PCC to the rest of the brain has shown to be altered in chronic pain and also in depression; depressive symptoms also have been shown to modulate connectivity disruptions of the PCC in chronic pain populations (23,29,59). Consistent with this literature, our connectivity analyses using PPI revealed that JFM patients exhibited hyperconnectivity between the PCC (including vPCC, Brainnetome area 23) and the left middle temporal gyrus (MTG), a posterior DMN region implicated in higher-order auditory and multimodal processing, sensory integration, memory, and emotion processing (60). Increased PCC–MTG connectivity in JFM may reflect altered functional integration within DMN-related networks, which are involved in dynamic interactions with salience and executive control systems (15,61). Such alterations may contribute to atypical processing of sensory and affective information even in the absence of nociceptive input (17,23). Specifically, the left MTG has been correlated with pain catastrophizing, rumination and affective disturbances in chronic pain and depressive disorders (62,63), and similar PCC-MTG hyperconnectivity has been reported in children with depression and link to emotion regulation difficulties (64). Taken together, these findings suggest a potential neurobiological vulnerability involving maladaptive emotion regulation and persistent self-referential processing during multisensory processing, supporting the hypothesis that affective symptoms may exacerbate sensory perception, including pain, in JFM.

Additionally, depressive symptoms (CDI-2 total score) were positively associated with increased connectivity between the PCC and a distributed set of regions including the hippocampus, thalamus, and cerebellum, suggesting alterations in network-level integration of memory, affective, and sensory processes. Notably, PCC–thalamus connectivity emerged as a central finding, involving predominantly thalamic subdivisions connected with the posterior parietal cortex, which are strongly implicated in the integration and gating of sensory information (65), and has been consistently altered in chronic pain conditions (17,19). This pattern may indicate that self-referential and affective processes more strongly influence early-stage sensory processing, contributing to the heightened salience of non-painful stimuli. In parallel, enhanced PCC–hippocampus connectivity may reflect altered encoding and emotional appraisal of pain-related experiences, consistent with evidence of corticolimbic dysfunction in chronic pain (66). Finally, increased PCC–cerebellar connectivity further supports the involvement of higher-order regulatory systems, given the cerebellum’s role in cognitive control, emotional regulation, and modulation of sensory processing (Zhang et al., 2023). Together, these findings point to distributed network-level alterations underlying the interaction between depressive symptoms and sensory hypersensitivity in JFM. Importantly, given the multimodal nature of the task (visual, auditory, and motor components), exploratory functional decoding of the unthresholded statistical maps revealed associations with terms related to visual processing, hand movements, and motor coordination, supporting the interpretation that PCC alterations emerge in the context of multisensory integration demands rather than isolated affective processing. This interpretation aligns with models of chronic pain emphasizing altered sensory integration and amplification across multiple modalities.

This study presents several limitations. First, only female participants were included, which limits the generalizability of our findings to male, transgender, or non-binary individuals. Future research should examine potential sex and gender differences in JFM. Second, despite efforts to recruit a diverse sample, the cohort lacked substantial ethnic and socioeconomic diversity. Community-engaged research approaches are essential to address the overrepresentation of white participants with medium-to-high socioeconomic status in clinical studies. Lastly, as this is the first study assessing depressive symptoms and its relation to non-painful sensory hypersensitivity in JFM using this approach, replication is necessary to confirm the robustness and translational relevance of our findings.

In conclusion, this study provides the first evidence of an association between sensory hypersensitivity in JFM and depressive symptoms and suggests that this relationship may be reflected in altered PCC-related neurofunctional processing. These findings highlight the interplay between sensory processing and emotional regulation in adolescents with JFM and provides novel insights into the early neurobiological mechanisms.

## Acknowledgments

This work was supported by Cincinnati Children’s Hospital Medical Center’s Trustee Grant Award (awarded to MLS) and NIH/NIAMS Grants R01 AR074795 and P30 AR076316, the PID2023-147704OB-I00 funded by MICIU/AEI/10.13039/501100011033/FEDER, UE granted to MLS and the CNS2023-14542 funded by MICIU/AEI/10.13039/501100011033 granted to MLS, NextGenerationEU/PRTR granted to MLS and AMIGO Project ‘A multi-ingredient brain function model predicting chronic pain in youth: a window into future well-being’, reference number 101171809, funded by the European Commission through the HORIZON EUROPE programme, European Research Council (ERC) Consolidator Grant granted to MLS. Marina López-Solà, PhD is hired as part of the Serra Hunter Programme of the Generalitat de Catalunya.

