## Supplementary Material for "Multisensory brain processing is associated with depressive symptoms in youth with chronic pain"

### Supplementary Methods and Materials

- **Supplementary Methods 1:** Recruitment Criteria
- **Supplementary Methods 2:** MRI acquisition parameters and Imaging processing
- **Supplementary Methods 3:** fMRI analysis
- **Supplementary Figure 1.** Percentage of overlap between PPI seed regions (PCC, ventral PCC subregion [Brainnetome area 23], and dorsal PCC subregion [Brainnetome area 31]) and brain responses during non-painful multisensory stimulation significantly associated with depressive symptoms (CDI-2 and CDI-2 subscales: Negative Mood, Interpersonal Problems, Anhedonia, Negative Self-Esteem, and Ineffectiveness).
- **Supplementary Figure 2.** Brain maps showing PPI connectivity differences for each PCC seed (v23ab, d23ab, 31pv, 31pd) between JFM patients and healthy controls and correlations between PPI-based connectivity maps for each seed and depressive symptoms in JFM patients.
- **Supplementary Table 1.** PCC subregions' hyperconnectivity in JFM patients compared to healthy controls and correlations with depressive symptoms (CDI-2 total) in JFM
- **Supplementary Table 2.** Correlations in whole brain between depressive symptoms and multisensory fMRI task
- **Supplementary Table 3.** PCC hyperconnectivity in JFM patients compared to healthy controls and correlations with depressive symptoms (CDI-2 total) in JFM
- **References of Supplementary Methods and Materials**

**Supplementary Methods 1: Participants and recruitment Criteria**

Participants were drawn from the Fibromyalgia Integrative Training program for Teens (FIT-teens) cohort (1). Juvenile fibromyalgia (JFM) was diagnosed according to established clinical criteria, and healthy controls were recruited from the same community and screened to exclude chronic pain conditions and major medical or psychiatric disorders. Detailed inclusion and exclusion criteria are provided in the original FIT-teens publication.

The sample was restricted to female adolescents due to the higher prevalence of JFM in girls and the composition of the parent clinical trial, in which 86% of eligible participants were cisgender girls. This limited the feasibility of including other genders and helped reduce potential confounding effects related to sex differences in pain processing and brain development (2–4).

JFM patients were recruited at the Cincinnati Children's Hospital Medical Center, after being diagnosed by a pediatric rheumatologist or a pain physician using 2010 American College of Rheumatology criteria. Healthy participants were recruited using the Office of Computational and Technical Research (OCTR) participant database of people interested in being notified about studies being conducted at Cincinnati Children's Hospital and from previous studies in which participants consented to be contacted for new research opportunities.

Inclusion criteria were the following: 1) A JFM diagnosis confirmed by a pediatric rheumatologist or pain physician using the 2010 American College of Rheumatology (ACR) criteria adapted for pediatric use. These criteria were chosen because they have been specifically adapted and validated in adolescent females with JFM and were also applied in the parent clinical trial FIT-teens (1). 2) An average pain intensity of 3 or greater out of 10 over the past week, as measured by the Brief Pain Inventory (BPI) (Daut et al., 1983), and 3) Functional Disability Inventory (FDI) scores higher than 7 out of 60, indicating at least mild disability (5). Inclusion criteria for healthy controls included 1) being physically and psychologically healthy (i.e., not diagnosed with chronic pain, psychiatric, neurological, or inflammatory disorders), and 2) average pain score of 0 on a Numeric Rating Scale and an FDI score equal or below 7.

Subjects with contraindication to MRI scanning, developmental delay, major neurological or psychiatric disorders (except for JFM patients with mood, anxiety, or obsessive-compulsive disorders undergoing treatment), a positive pregnancy test, or taking opioid medication were not eligible. All participants included in the study were under no medication or under a stable medication regime for a minimum of 3 weeks prior to the MRI assessment.

All participants and their parent/legal guardians provided written informed consent/assent. The Cincinnati Children's Hospital Medical Center's Institutional Review Board approved the study protocol and consent forms (ID: 2017-7771).

**Supplementary Methods 2: MRI acquisition parameters and Imaging processing**

We scanned participants with a Philips Ingenia 3.0-Tesla MRI System (Philips Healthcare, Best, Netherlands) equipped with a 32-channel head coil at the Cincinnati Children's Hospital Medical Center using a T2* weighted echo planar imaging sequence with a multiband sensitivity encoding (SENSE) technique and the following scan parameters: multiband acceleration factor = 4, TR = 650 ms, TE = 30 ms, field of view = 200 mm, flip angle = 53°, voxel size =2.5 x 2.5 x 3.5 mm, slice orientation = transverse (parallel to the orbitofrontal cortex line), slice thickness = 3.5 mm, number of slices = 40 (providing whole-brain coverage), number of volumes = 433, dummy scans = 12, and total scan duration = 4,69 min. T1-weighted structural images were reviewed by a board-certified neuroradiologist. No clinically significant anatomical abnormalities were detected.

Imaging data were processed using gold standard preprocessing pipelines implemented in the CONN Toolbox version 21.a (www.nitrc.org/projects/conn) running on MatLab version 2021a (Math Works Inc, Natick, MA). Standard preprocessing steps included realignment and unwarping, outlier detection, spatial normalization to MNI, and smoothing. The first 12 volumes of the multisensory sequence were discarded to allow magnetization to reach equilibrium, and the remaining volumes were realigned with the first one. Based on head-motion parameters estimated during realignment, volumes with framewise displacement >0.9mm or global BOLD signal changes >5 standard deviations were flagged as outliers. Subsequently, fMRI data were normalized to the 2mm isotropic MNI space using direct functional normalization, which estimates tissue types and computes the nonlinear transformation directly from the mean BOLD signal image, rather than relying on the anatomical scan (6,7). This approach was chosen both because one participant lacked a T1-weighted image and because prior evidence shows that direct functional normalization reduces inter-subject variability, improves alignment, and increases activation sensitivity (8–10). Last, functional images were smoothed with an 8mm full width at half-maximum Gaussian kernel to optimize detection of regions with large spatial extents, previously shown to be evoked during the multisensory task. Images were inspected for artifacts at every step and quality assurance plots were visually examined. No participants were excluded based on motion, as none exceeded predefined thresholds warranting subject-level exclusion.

**Supplementary Methods 3: fMRI analysis**

In first-level analyses, we computed generalized linear models in SPM12 (The Wellcome Department of Imaging Neuroscience) for each participant to identify brain regions with significant activation changes during the multisensory blocks compared to the baseline condition. A primary task regressor was created by convolving the stimulation blocks with a canonical hemodynamic response function. Similarly, we also modeled the unpleasantness question that followed each trial as a regressor. The rest condition served as the implicit task baseline. A high-pass filter was used to remove low-frequency signal fluctuations (1/180 Hz). Average framewise displacement and outlier volumes (i.e., volumes with framewise displacement >0.9mm or global BOLD signal changes >5 standard deviations, flagged as outliers during data preprocessing) were regressed out in subject-level analyses.

Contrast images for the task regressor of interest obtained for each subject were carried forward to second level group analyses. Contrast images for the task regressor of interest for each subject were carried forward into separate group regression models to compute the parametric association between brain activation and depressive symptoms (CDI-2). Statistical significance was set at family-wise error (FWE) $p_{FWE-}<0.05$ at the cluster level and $p<0.001$ at the voxel level. As post-hoc analysis we wanted to see the correlation between brain activation and the different CDI-2 subscales (Negative Mood, Interpersonal Problems, Negative Self-Esteem, and Ineffectiveness). We found no significant differences at a whole-brain FWE cluster-corrected level of $p<0.05$ with voxel-level $p<0.001$ in the Interpersonal Problem and the Negative Self-Esteem subscales. We additionally report exploratory results for this subscale at a more lenient threshold of $p<0.005$ (cluster extent >20 voxels), uncorrected, consistent with common practice in neuroimaging, which should be interpreted as hypothesis-generating .

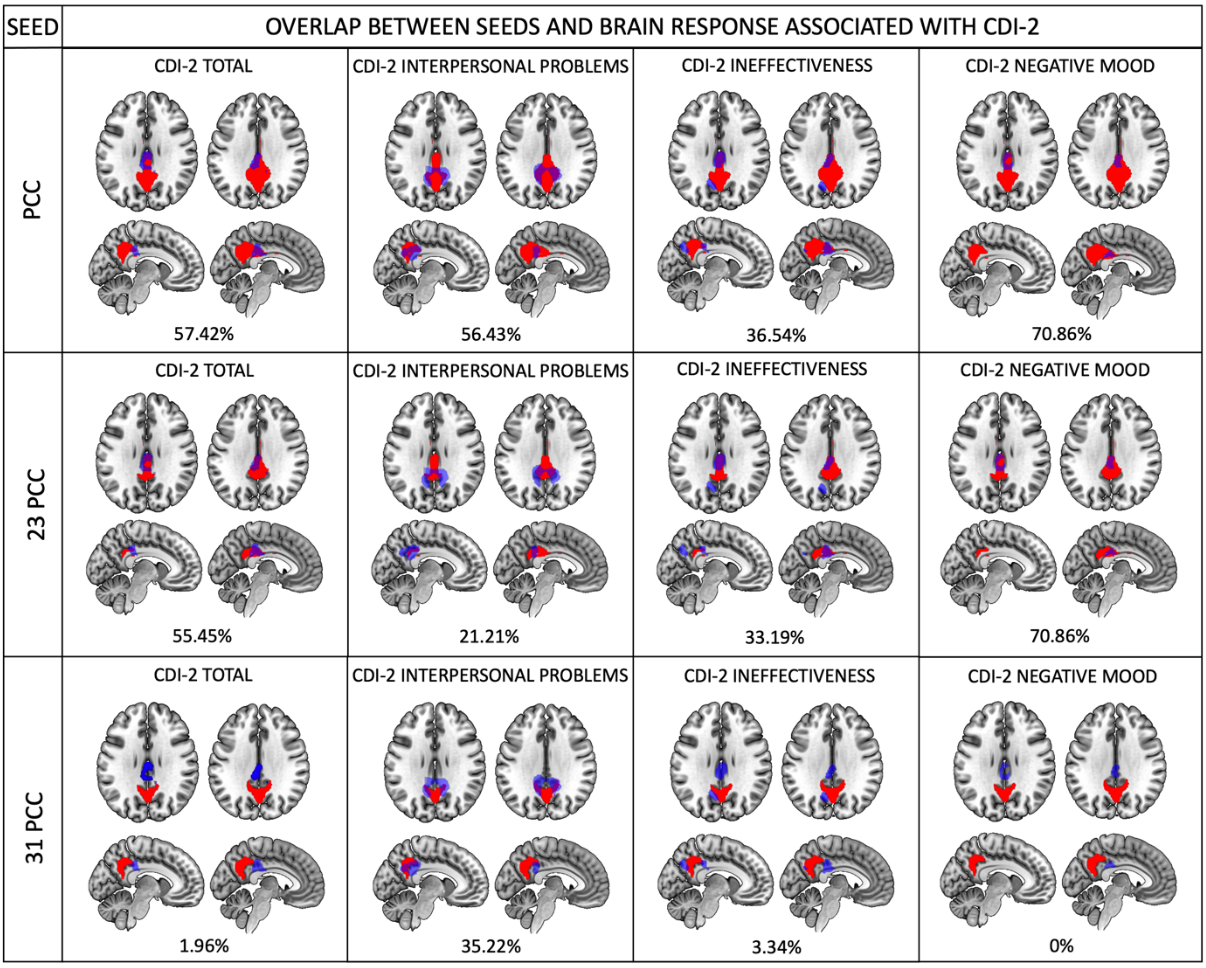


**Supplementary Figure 1. Percentage of overlap between PPI seed regions (PCC, ventral PCC subregion [Brainnetome area 23], and dorsal PCC subregion [Brainnetome area 31]) and brain responses during non-painful multisensory stimulation significantly associated with depressive symptoms (CDI-2 and CDI-2 subscales: Negative Mood, Interpersonal Problems, Anhedonia, Negative Self-Esteem, and Ineffectiveness).** The color red represents the PPI seeds, the color blue represents the brain response activation associated with depressive symptoms and the color purple is the overlap between both regions. The percentage represents the overlap between the clusters of the two regions.

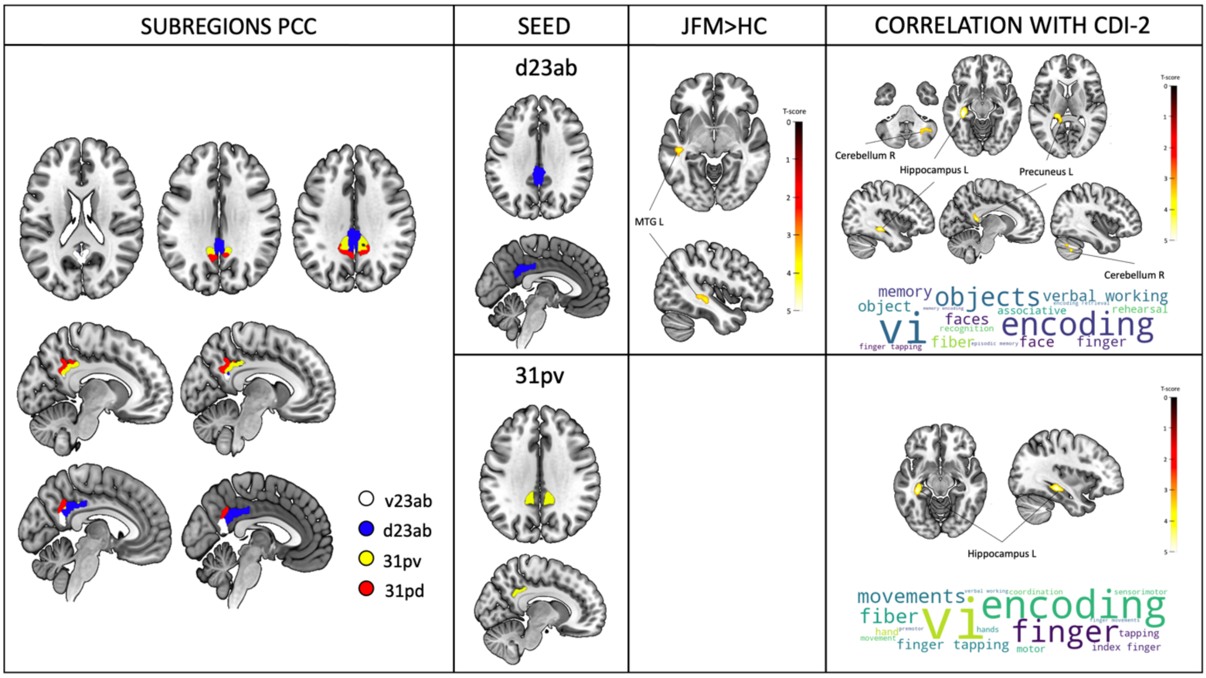


**Supplementary Figure 2. Brain maps showing PPI connectivity differences for each PCC seed (v23ab, d23ab, 31pv, 31pd) between JFM patients and healthy controls and correlations between PPI-based connectivity maps for each seed and depressive symptoms in JFM patients.** The bottom right of the figures shows the 15 functional terms most associated with the unthresholded brain t-map of results for each analysis based on the meta-analytic database of Neurosynth (11). Wordcloud was computed with Python. The size of the word is based on the correlation coefficient between each specific term and the connectivity t-map. The color of words is for aesthetic purposes.

**Supplementary Table 1.** PCC subregions' hyperconnectivity in JFM patients compared to healthy controls and correlations with depressive symptoms (CDI-2 total) in JFM

| | Cluster size (voxels) | *MNI coord.* x y z | z | pFWE | Cohen's D |
|---|---|---|---|---|---|
| **JFM<HC** | | | | | |
| **d23ab PCC** | | | | | |
| Middle Temporal Gyrus Left | 172 | -44 -26 -6 | 3.91 | 0.012 | 0.86 |
| **CORRELATIONS WITH CDI-2 TOTAL** | | | | | |
| **d23ab PCC** | | | | | |
| Caudal Hippocampal Left | 197 | -34 -32 -12 | 4.79 | 0.009 | 1.74 |
| Thalamus Left | 188 | -10 -44 12 | 4.22 | 0.011 | 1.48 |
| Cerebellum Right | 169 | 36 -58 -44 | 3.85 | 0.019 | 1.33 |
| **31pv PCC** | | | | | |
| Caudal Hippocampal Left | 164 | -36 -30 -12 | 4.50 | 0.022 | 1.61 |

Note: Cluster size is presented in number of voxels. JFM, Juvenile fibromyalgia; MNI coord, Montreal Neurological Institute coordinates; PCC, posterior cingulate cortex.

**Supplementary Table 2.** **C**orrelations between depressive symptoms and whole brain activation during the non-painful multisensory fMRI task

| | Cluster size (voxels) | *MNI coord.* x y z | z | pFWE | Cohen's D |
|---|---|---|---|---|---|
| **DEPRESSIVE SYMPTOMS** | | | | | |
| **CDI-2 Total** | | | | | |
| Right Dorsal PCC | 1345 | 0 -28 42 | 3.95 | 0.002 | 1.37 |
| **CDI-2 Interpersonal Problems** | | | | | |
| Right Ventral PCC | 1169 | 10 -44 32 | 4.02 | 0.005 | 1.39 |
| **CDI-2 Ineffectiveness** | | | | | |
| Right Dorsal PCC | 1827 | 4 -28 24 | 4.32 | <0.001 | 1.52 |
| **CDI-2 Negative Mood** | | | | | |
| Right Dorsal PCC | 821 | 2 -28 42 | 3.94 | 0.028 | 1.36 |

Note: Cluster size is presented in number of voxels. CDI-2, Child Depression Inventory 2; MNI coord, Montreal Neurological Institute coordinates; PCC, posterior cingulate cortex.

**Supplementary Table 3.** PCC hyperconnectivity in JFM patients compared to healthy controls and correlations with depressive symptoms (CDI-2 total) in JFM

| | Cluster size (voxels) | *MNI coord.* x y z | z | pFWE | Cohen's D |
|---|---|---|---|---|---|
| **JFM<HC** | | | | | |
| **PCC Total** | | | | | |
| Middle Temporal Gyrus Left | 306 | -44 -26 -6 | 4.14 | <0.001 | 0.92 |
| **23 PCC** | | | | | |
| Middle Temporal Gyrus Left | 172 | -44 -26 -6 | 3.91 | 0.012 | 0.87 |
| **CORRELATIONS WITH CDI-2 TOTAL (in JFM)** | | | | | |
| **PCC Total** | | | | | |
| Caudal Hippocampal Left | 187 | -34 -34 -10 | 4.49 | 0.011 | 1.60 |
| • Rostral Ventral Area 23 | 4 | | | | |
| • Posterior Parahippocampal gyrus | 1 | | | | |
| • Rostral hippocampal | 15 | | | | |
| • Caudal hippocampus | 90 | | | | |
| Thalamus Left | 171 | -10 -44 12 | 4.26 | 0.017 | 1.50 |
| • Caudal hippocampus | 7 | | | | |
| • Rostral Temporal Thalamus | 8 | | | | |
| • Posterior Parietal Thalamus | 27 | | | | |
| • Occipital Thalamus | 11 | | | | |
| • Caudal Temporal Thalamus | 23 | | | | |

| | | | | | |
|---|---|---|---|---|---|
| Cerebellum Right | 375 | 38 -66 -36 | 4.19 | <0.001 | 1.47 |
| **23 PCC** | | | | | |
| Caudal Hippocampal Left | 229 | -34 -32 -12 | 4.84 | 0.004 | 1.76 |
| Caudal Temporal Thalamus Left | 286 | -20 -32 12 | 4.71 | 0.001 | 1.70 |
| Cerebellum Right | 327 | 38 -66 -36 | 3.95 | <0.001 | 1.37 |
| **31 PCC** | | | | | |
| Cerebellum Right | 204 | 38 -66 -36 | 4.03 | 0.008 | 1.40 |

Note: Cluster size is presented in number of voxels. JFM, Juvenile fibromyalgia; MNI coord, Montreal Neurological Institute coordinates; PCC, posterior cingulate cortex.
